# Impact of non-LTR retrotransposons in the differentiation and evolution of Anatomically Modern Humans

**DOI:** 10.1101/207241

**Authors:** Etienne Guichard, Valentina Peona, Guidantonio Malagoli-Tagliazucchi, Lucia Abitante, Evelyn Jagoda, Margherita Musella, Marco Ricci, Alejandro Rubio-Roldán, Stefania Sarno, Donata Luiselli, Davide Pettener, Cristian Taccioli, Luca Pagani, Jose Luis Garcia-Perez, Alessio Boattini

## Abstract

Transposable Elements are biologically important components of eukaryote genomes. In particular, non-LTR retrotransposons (N-LTRrs) extensively shaped the human genome throughout evolution. In this study, we compared retrotransposon insertions differentially present in the genomes of Anatomically Modern Humans, Neanderthals, Denisovans and Chimpanzees, in order to assess the possible impact of retrotransposition in the differentiation of the human lineage. Briefly, we first identified species-specific N-LTRrs and established their distribution in present day human populations. These analyses shortlisted a group of N-LTRr insertions that were found exclusively in Anatomically Modern Humans. Notably, these insertions targeted genes more frequently than randomly expected and are associated with an increase in the number of transcriptional/splicing variants of those genes they inserted in. The analysis of the functionality of genes targeted by human-specific N-LTRr insertions seems to reflect phenotypic changes that occurred during human evolution. Furthermore, the expression of genes containing the most recent N-LTRr insertions is enriched in the brain, especially in undifferentiated neurons, and these genes associate in networks related to neuron maturation and migration. Additionally, we also identified candidate N-LTRr insertions that have likely produced new functional variants exclusive to modern humans, which show traces of positive selection and are now fixed in all present-day human populations. In sum, our results strongly suggest that N-LTRr impacted our differentiation as a species and have been a constant source of genomic variability all throughout the evolution of the human lineage.

## INTRODUCTION

Transposable Elements (TEs) are DNA sequences that are able to move or replicate in genomes via cut-and-paste and copy-and-paste-like mechanisms (Richardson et al. 2015). Although TEs have for long been dismissed as "selfish", "parasites" or simply "junk DNA" (Orgel and Crick, 1980; Doolittle and Sapienza, 1980), the advent of whole genome DNA sequencing, in conjunction with molecular genetic, biochemical, genomic and functional studies, has revealed that TEs are biologically important components of mammalian genomes whose activity has extensively shaped the structure and function of our own genome (Richardson et al. 2015).

TEs are known to be involved in several evolutionary and adaptive processes such as the generation of genes and pseudogenes (Ohshima et al. 2003; Moran et al. 1999; Sayah et al. 2004), fine-tuning transcriptional regulation of genes (Speek 2001; Han et al. 2004; Chuong et al. 2017), generation of somatic mosaicism (Bailie et.al. 2011; Muotri et al. 2005; Evrony et al. 2012), the increase in complexity and evolution of gene regulatory networks (Feschotte 2008) and the alteration of epigenetic mechanisms as processes of fine-scale and reversible regulation (Fedoroff 2012). Some of the most notable biological processes associated with the domestication of TE-derived sequences are the insurgence of the V(D)J system of acquired immunity (Kapitonov and Jurka, 2005; Koonin and Krupovic, 2014; Huang et al. 2016) and placental development (Lynch et al. 2011, Lavialle et al. 2013), but they also play key roles in embryogenesis (Gerdes et al. 2016; Friedli and Trono, 2015) and neurogenesis (Notwell et al. 2015; Muotri et al. 2005; Evrony et al. 2012). In sum, in addition to their role in growing the size of eukaryotes' genomes, active TEs are continually impacting the functioning and evolution of genomes.

Notably, the activity of TEs throughout evolution has generated more than two thirds of the human genomic material (Lander et al. 2001; Kapusta et al. 2013). In modern humans, only a limited number of TE subfamilies from the non-LTR-retrotransposon (N-LTRr) class are currently active, i.e. Long and Short Interspersed elements (LINE-1s and SINEs, including Alus and SVAs). Indeed, the ongoing activity of LINE-1s and SINEs in humans offers a constant source of inter-individual variability in human populations and can sporadically generate new genetic disorders (Kazazian et al. 1988; Richardson et al. 2015).

In this study, we aimed to explore the role of N-LTRrs in the differentiation and evolution of the genus *Homo*. In order to do that, we compared the repertoire of Retrotransposon Insertions (RI) present in the genome of modern humans with those present in modern-day chimpanzees, as well as with those of our closest extinct relatives, Neanderthals and Denisovans. *Homo neanderthalensis* (HN) and *Denisova* (HD) are sister groups, being more closely related to each other than they are to *Homo sapiens*. Their split from the modern human lineage is estimated to have occurred between 550 thousands of years ago (Kya) and 765 Kya, after which they colonized Eurasia long before Anatomically Modern Humans (AMH) left Africa. The population split between these archaic populations is estimated at 381-473 kya (Prüfer et al. 2014). Notably, the genomes of some individuals belonging to HN and HD have been previously sequenced, assembled and published (Green et al. 2010; Meyer et al. 2012; Prüfer et al. 2014; Sawyer et al. 2015).

Although often discussed and speculated, the effects and implications of RIs on the evolution of the human lineage are mostly unknown. Here, and for the first time, we evaluated the potential impact of RIs in our species differentiation; additionally, we have also analyzed how RIs that are specific to AMH and absent in HN, HD and chimpanzees could have affected the genomic loci surrounding them. In order to shed light on the molecular dynamics of AMH differentiation and evolution, we aimed at characterizing Rl locations, identifying potential selective pressures and inferring functional/regulatory alterations that might have occurred as a consequence of species-specific RIs. Thus, the reconstruction of the mechanisms through which retrotransposition impacted the evolution of the human lineage can allow for a better understanding of how our genome is evolving in real time.

## RESULTS

### RI identification

Available RI identification tools such as RetroSeq (Keane et al. 2012), Tangram (Wu et al. 2014), Tea (Lee et al. 2012), MELT (Gardner et al. 2017), etc. are primarily based on mapping paired-end DNA-sequencing reads. However, and given that a large portion of previously sequenced ancient DNAs is composed of single-ended reads, here we devised a methodology for detecting differentially present RIs in AMH, HD and HN genomes based on single-ended reads (Supplemental Figure S1). In particular, our methodology (see METHODS) is intended to identify confirmed species-specific RIs upon which to infer the impact that RIs might have had in our species differentiation.

These analyses led to the identification of: i) 507 HN-specific and 331 HD-specific putative RIs, and ii) 3215 and 7185 putative AMH-specific RIs vs HN and HD, respectively.

As for the comparison between AMH (GRCh37-hg19) and chimpanzee (panTro5) genomes, we retrieved all RIs annotated in RepBase (Bao et al. 2015) in these two genomes and analyzed the presence/absence of the insertions in the reference sequences of the two species.

Next, we developed a computational validation procedure, through which we managed to eliminate all those insertions that presented uncertainties in mobile element subfamily attribution or whose location might be ambiguous (see METHODS for details). Thus, we only continued the analysis of the most reliable canonically-inserted RIs identified (Supplemental Figure S1).

A number of species-specific RI were computationally validated: 1370 Chimp-specific, 38 HD-specific, 64 HN-specific, 5402 AMH-specific (against chimps), 548 AMH-specific (against Denisova) and 806 AMH-specific (against Neanderthal) (Table 1, Supplemental Tables S1-S7). The validation method thus excluded approximately 87% of the identified insertions that could present any sort of bias or uncertainty in attribution. Of the validated AMH-specific RIs, 321 were present in the modern human genome and were absent in both HN and HD genomes.

**Table 1.** Identified and validated RIs in chimpanzee, HN, HD and AMH genomes.

|  | **TOTAL** | **Alu** | **LINE-1** | **SVA** |
| --- | --- | --- | --- | --- |
| **Chimp-specific RIs** | 1906 | 1370 | 463 | 73 |
| **HN-specific RIs** | 64 | 57 | 6 | 1 |
| **HD-specific RIs** | 38 | 32 | 6 | 0 |
| **AMH-specific RIs vs chimp** | 5402 | 4504 | 655 | 253 |
| **AMH-specific RIs vs HN** | 806 | 728 | 77 | 1 |
| **AMH-specific RIs vs HD** | 548 | 487 | 58 | 3 |
| **AMH-specific RIs vs both HN and HD** | 321 | 295 | 25 | 1 |

The comparison of the identified insertions with a large dataset of ^~^666,000 retrotransposon insertions from the most recent subfamilies of N-LTRrs (i.e, Alus, LINE-1s and SVAs) present in the reference GRCh37-hg19 (defined as RT-DB) revealed that the activity of N-LTRrs in the human lineage has remained constant. Consistently, Alu RIs are far more common than LINE-1 RIs, while SVAs produced only a handful of insertions. These results strongly suggest that the retrotransposition rate, or insertion maintenance rate, in the human lineage has remained relatively constant (0.6-0.8 insertions/Ky), and that the rate of RI accumulation in humans has been approximately 2.5 times higher than in chimps (0.29 insertions/Ky).

### Archaic-specific RIs and insertional polymorphisms

HD-and HN-specific RIs (38 and 64, respectively) were compared between the two species and with present-day AMH populations data from 1000 Genomes Project Phase 3 (Supplemental Tables S8 and S9, Supplemental Figure S2A-B) (The 1000 Genomes Project Consortium 2015). Based on the available 1000 Genomes Project data, three RIs were found in both archaic species, while nearly half of them (49 out of 102) are polymorphic to various degrees in modern populations. Interestingly, 8 of the insertions (1 HD-specific and 7 HN-specific) are absent in African (AFR) individuals and present at a low frequency only in some (or all) non-AFR populations. Thus, we speculate that these RIs might have introgressed in AMH via admixture with the archaic species after *Homo sapiens* migrated out of Africa. Conversely, given the documented presence of Neanderthal introgressed sequences within the genome of Eurasians (Green et al. 2010; Prüfer et al. 2014; Gardner et al. 2017), which in turn forms the majority of the human reference sequence, it may be the case that HN and HD specific insertions present on the human reference due to archaic introgression escaped our detection.

Since some putative archaic-specific RIs are polymorphic in modern humans, it is likely that at least some putative modern-specific insertions might be polymorphic in archaic populations as well; however, we would need many more available ancient genomes to test this. In order to estimate the number of potential polymorphic AMH-specific insertions, we took advantage of the large amount of population genetics data provided by the 1000 Genomes project. Indeed, by randomly sampling AFR individuals we observed that a few samples would be sufficient to identify the vast majority of the archaic-specific polymorphic insertions, reaching a plateau at n=20. Similarly, ^~^45% of the putative species-specific insertions were shown to be polymorphic (Supplemental Figure S2C). On the other hand, in the AMH-specific insertions, we observed that the 321 detected RIs (100%) that were present in AMH and absent in both archaic genomes fall below the ~45% threshold identified with the above procedure. This fact, together with the observation that HD and HN genomes are more divergent than two randomly-chosen AMH genomes (Reich et al. 2010), suggests that the above mentioned 321 insertions may be considered as reliable and truly AMH-specific.

### AMH-specific RIs in present-day populations

The fact that the detected 321 AMH-specific RI are present in the human reference sequence (GRCh37-hg19) does not necessary imply that they are fixed in all human populations. We therefore evaluated their distribution in present-day populations by comparing the coverage of the unique 3’ and 5’ flanking regions with that of the RI/flanking interface in 1000 Genomes Phase 3 data (Supplemental Table S10, Supplemental Figure S2D; more details in METHODS). This analysis revealed that, of the 321 AMH-specific insertions, 24 (7,5%) appear to be fixed or almost fixed in all modern populations (allele frequency > 85%), while 8 (2,5%) are polymorphic in AFR individuals but fixed or almost fixed in all non-African populations (allele frequency < 65% in AFR and > 85% in non-Africans), suggesting that their fixation may be related to the Out-of-Africa event.

Interestingly, the patterns of RI distribution seem to closely reflect known pre-historic and historic migrations and population dynamics of AMH (Supplemental Figure S2E). In particular, populations of African descent are the more divergent and the Out-of-Africa groups cluster according to clear phylogenetic/phylogeographic relationships, with the expected exceptions of PUR and CLM who cluster with EUR populations and not with AMR, likely because of admixture during the re-colonization of North and South America (Montinaro et al. 2015).

Time to the Most Recent Common Ancestor (TMRCA) were also calculated for 10kbp sequences (Inchley et al. 2016) surrounding each insertion site (Supplemental Table S10, details in METHODS). Of the 24 insertions that are fixed or almost fixed in al modern populations, we selected those showing a TMRCA compatible with the split between AMH and HN/HD (TMRCA < 800 Kya) as potential candidates for selection/spread along the AMH lineage. Accordingly, we identified two RIs (8%), i.e. an AluYg6 insertion in chr1q25.3 that occurred in the gene EDEM3 and an AluYb9 insertion in chr10q25.3 that also occurred within the sequence of the gene SHTN1. Similarly, only one of the 8 AMH-specific insertions that are likely fixated post Out-of-Africa, an AluYa5 insertion in chr16q22.1, also displays a recent TMRCA. However, it is worth noting that TMRCA estimates were obtained from all the AFR individuals and not only from the carriers of an insertion; therefore, they are therefore to be considered as a general indicator of the “age” of a given site surrounding an insertion or, in other words, as an upper-limit for the retrotransposition event itself.

Three Population Composite Likelihood Ratio (3P-CLR) statistic (Racimo 2016) was also performed on 200kbp loci surrounding each insertion. This analysis revealed that 28 (9.2%) out of the 306 AMH-specific RIs autosomal insertional loci are within the top 0.1% of loci subjected to post Out-of-Africa selection (Supplemental Table S11, details in METHODS).

### Genomic features of loci targeted by AMH-specific RIs

The huge amount of genetic and genomic data presently available on modern humans allowed us to perform different exploratory analyses on the AMH-specific RIs and their surrounding genomic loci, aimed at evaluating the possible impact of RIs in our species evolution.

First, we compared selected datasets of RIs (RT-DB, AMH-specific vs chimp and AMH-specific vs both archaic species) with the ENSEMBL gene annotation (Aken et al. 2016) of the reference sequence GRCh37-hg19. We determined that 15367 genes contain insertions of RT-DB (48.7% of the insertions), 1779 genes contain AMH-specific vs chimp RIs (43.9% of the insertions) and AMH-specific RIs targeted 139 genes after the split with HN/HD (43.3% of the insertions) (Figure 1A). These data suggests that RT-DB insertions targeted genes and gene-related sequences, or have been maintained throughout evolution in those sequences, ^~^30% more frequently than randomly expected in respect to gene-size/genome-size (p-value < 10^−16^), while AMH-specific RIs, both vs chimp and vs HN/HD, occurred in genes ^~^17% more frequently than expected (p-values < 10^−16^ and < 0.05 respectively).

**Figure 1.**
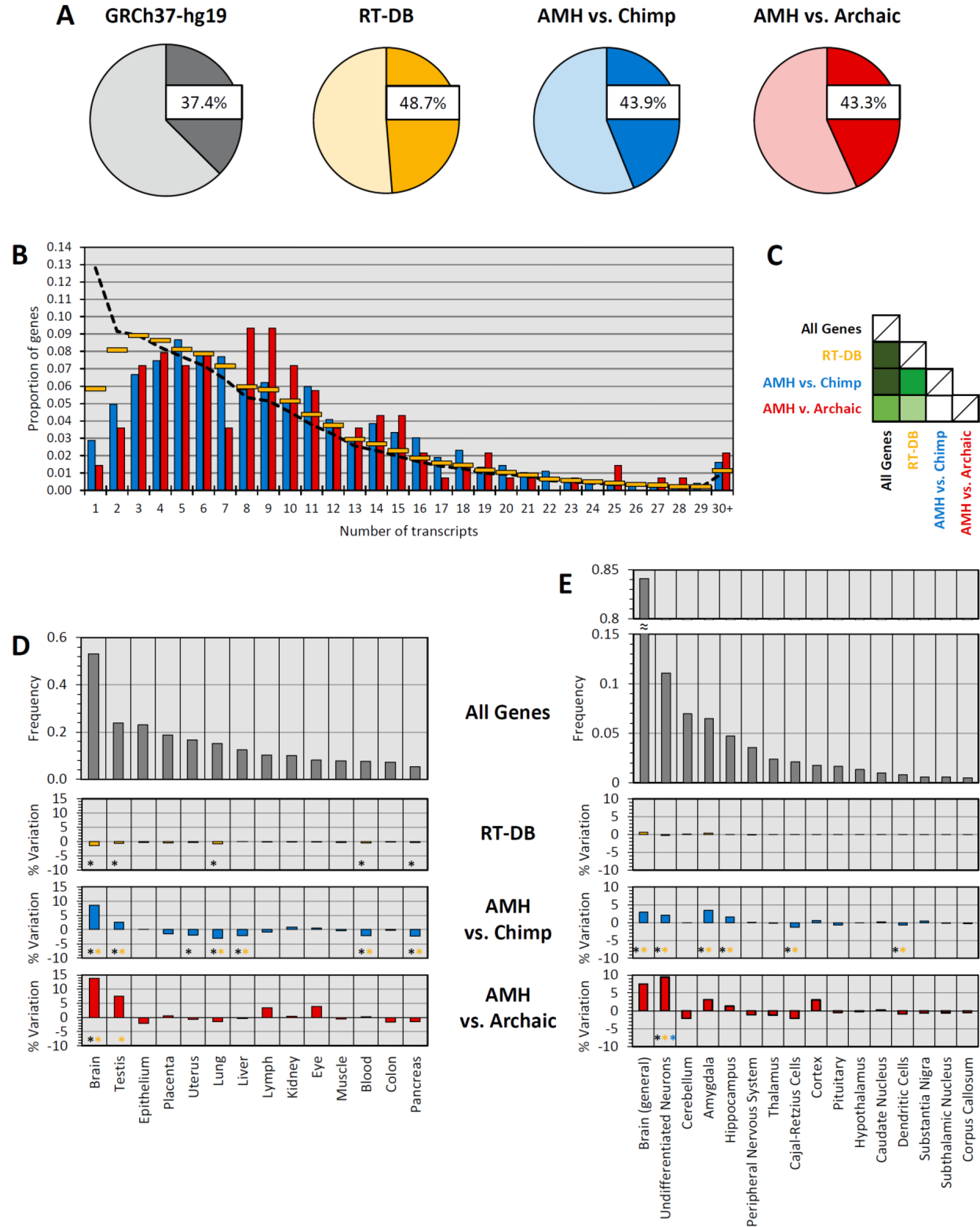
AMH-specific RI, genes and preferential expression. A) Proportion of ENSEMBL-annotated genes in the whole reference genome GRCh37-hg19 (grey), proportion of insertions that occurred in annotated genes for RT-DB insertions (yellow), AMH-specific RI vs chimp (blue) and AMH-specific RI vs both HN and HD (red). In each diagram, the darker color denotes the percentage of RIs inserted in genes vs RIs inserted in non-genic regions (lighter color). B) Proportion of genes per number of annotated transcripts for all ENSEMBL-annotated genes in the reference genome GRCh37-hg19 (black dotted line), for genes targeted by RT-DB insertions (yellow lines), for genes containing AMH-specific RIs vs chimp (blue bars) and for genes with AMH-specific RIs vs both HN and HD (red bars). C) Table showing statistical significance of the differences between the series of Figure 3B, calculated with Kolmogorov-Smirnov tests; white is for non-significant p-value, light-green is for p-value < 10^−2^, pea-green is for p-value < 10^−5^, emerald-green is for p-value < 10^−10^, dark-green is for p-values < 10^−16^. D) Proportion of all human genes showing preferential expression in different tissues (grey bars); % increase or decrease in absolute proportions for preferential tissue expression of genes targeted by RT-DB insertions (yellow bars), genes containing AMH-specific RIs vs chimp (blue bars) and genes with AMH-specific RIs vs both HN and HD (red bars). Black asterisks mark significant differences between the series and all human genes while yellow asterisks mark significant differences between the series and genes targeted by RT-DB insertions. E) Proportion of all human genes showing preferential expression in the brain divided by neural regions (grey bars); % increase or decrease in absolute proportions for preferential neural expression of genes targeted by RT-DB insertions (yellow bars), genes containing AMH-specific RIs vs chimp (blue bars) and genes with AMH-specific RIs vs both HN and HD (red bars). Black asterisks mark significant differences between the series and all human genes, yellow asterisks mark significant differences between the series and genes targeted by RT-DB insertions, blue asterisks mark significant differences between the series and genes targeted by AMH-specific RIs vs chimp.

In addition, the ENSEMBL gene-annotation data revealed that, in general, the majority of genes in AMH genomes tend to have a low number of annotated transcript/splicing variants, with a decreasing trend between the proportion of genes and the number of transcripts (7.584 on average; mode: 1 transcript/gene). Intriguingly, the comparison of genes targeted by RIs in the human lineage with all others present in AMH genomes (Figure 1B-C) revealed an average increase in the number of transcript and splicing variants for those genes that contain RT-DB insertions (8.428 on average, mode: 3 transcripts/gene; p-value < 10^−16^). Notably, this trend increases further when analyzing RIs that likely inserted after the split with chimps (9.728 on average, mode: 5 transcripts/gene; p-value < 10^−16^) and after the split with HN/HD (9.863 on average, mode: 8.5 transcripts/gene; p-value < 10^−6^). Consistently, genes targeted by AMH-specific RIs, both vs Chimp and vs HN/HD, also have more annotated transcripts than genes containing RT-DB insertions (p-values < 10^−11^ and < 0.005 respectively).

Next, we retrieved functional annotation data from DAVID Bioinformatics Resources v6.8 (Huang et al. 2009) and we obtained tissue-specific preferential gene expression information for 15126 out of 15367 genes with insertions from RT-DB, 1721 out of 1779 genes targeted by retrotransposition in the human lineage after the split with chimps and 124 out of 139 targeted after the split with HN/HD. Comparisons among these data show that genes targeted by retrotransposition tend to be more expressed than others in specific tissues (Figure 1D). Genes containing RT-DB insertions tend to follow the general expression profile of all human genes, with a slight under-expression in some tissues (p-values < 0.05); however, genes targeted by AMH-specific RIs after the split with chimpanzees are more expressed than average in the brain and testis (+8.5% and +2.7% in absolute proportions respectively; p-value < 10^−12^ and < 10^−2^), while being less expressed in the uterus, lungs, liver, blood and pancreas (decreases between −1.8% and −3.1% in absolute proportions; all p-values < 10^−2^); finally, genes with AMH-specific RIs absent in both HN and HD are significantly more expressed in the brain and, with respect to genes targeted by RT-DB insertions, in testis (+13.8% and +7.5% in absolute proportions, p-values < 5×10^−3^ and < 0.05).

We next analyzed genes preferentially expressed in the brain (Figure 1E) and observed that genes targeted by RT-DB insertions follow the same expression pattern of all human genes; genes with AMH-specific vs chimps RI are generally highly expressed in the brain and seem to be even more expressed than average in the amygdala and hippocampus, as well as in undifferentiated neurons (+3.4%, +1.5% and +2.0% in absolute proportions respectively, p-values < 10^−4^ for the amygdala and < 0.05 for both hippocampus and undifferentiated neurons), while showing less expression in Cajal-Retzius and dendritic cells (−1.4% and −0.7% in absolute proportions, p-values < 10^−3^ and < 0,05); finally, genes containing AMH-specific RIs absent in both HN and HD are significantly more expressed than average in undifferentiated neurons (+9.4% in absolute proportions, p-value < 10^−2^).

### Gene Ontology of genes targeted by AMH-specific RI

In order to examine the functionality of genes targeted by insertions, Gene Ontology (GO) analyses were performed on all genes targeted by AMH-specific RIs, both vs Chimp and vs HN/HD. ToppCluster analyses (Kaimal et al. 2010) revealed that, of the 238 GO terms identified between the two lists, 175 GO terms (73,5%) were overrepresented in genes targeted by AMH-specific RIs vs chimp, whereas 23 (9,7%) were overrepresented in genes targeted by AMH-specific RIs vs both HN/HD (Figure 2A, Supplemental Table S12). Next, we selected the GO terms that were enriched in one group of genes and not in the other as lineage-specific functionalities that might correspond to different moments in the evolution of the human lineage, i.e. hominid-specific GO terms (for terms enriched only in genes targeted by AMH-specific RI vs chimp) and sapiens-specific GO terms (for terms enriched only in genes targeted by AMH-specific RI vs both HN/HD). Interestingly, semantic similarity of hominid-specific GO terms showed that the most enriched functionalities of genes containing AMH-specific RI vs chimp are related to cognition, learning and memory capabilities, vocalization behavior, neuron recognition, dendrite morphogenesis, reflexes and regulation of locomotion (Figure 2B). Remarkably, these functionalities associate in networks involving a large number of genes containing AMH-specific RIs vs chimp. Even more noticeably, for enriched sapiens-specific GO terms, all functionalities associated in networks are neural-related: synapse maturation and its regulation, neuron maturation and migration, gliogenesis and glia differentiation (Figures 2C-D). Genes associated with these GO terms also form a complex network of interactions (Figure 2E). Strikingly, two of the genes with the larger amount of interactions in this network are SHTN1 and EDEM3 (1^st^ and 11^th^ scores in order of significance), which contain the previously identified (see above) AluYg6 RI (chr1q25.3) and the AluYb9 RI (chr10q25.3) respectively.

**Figure 2.**
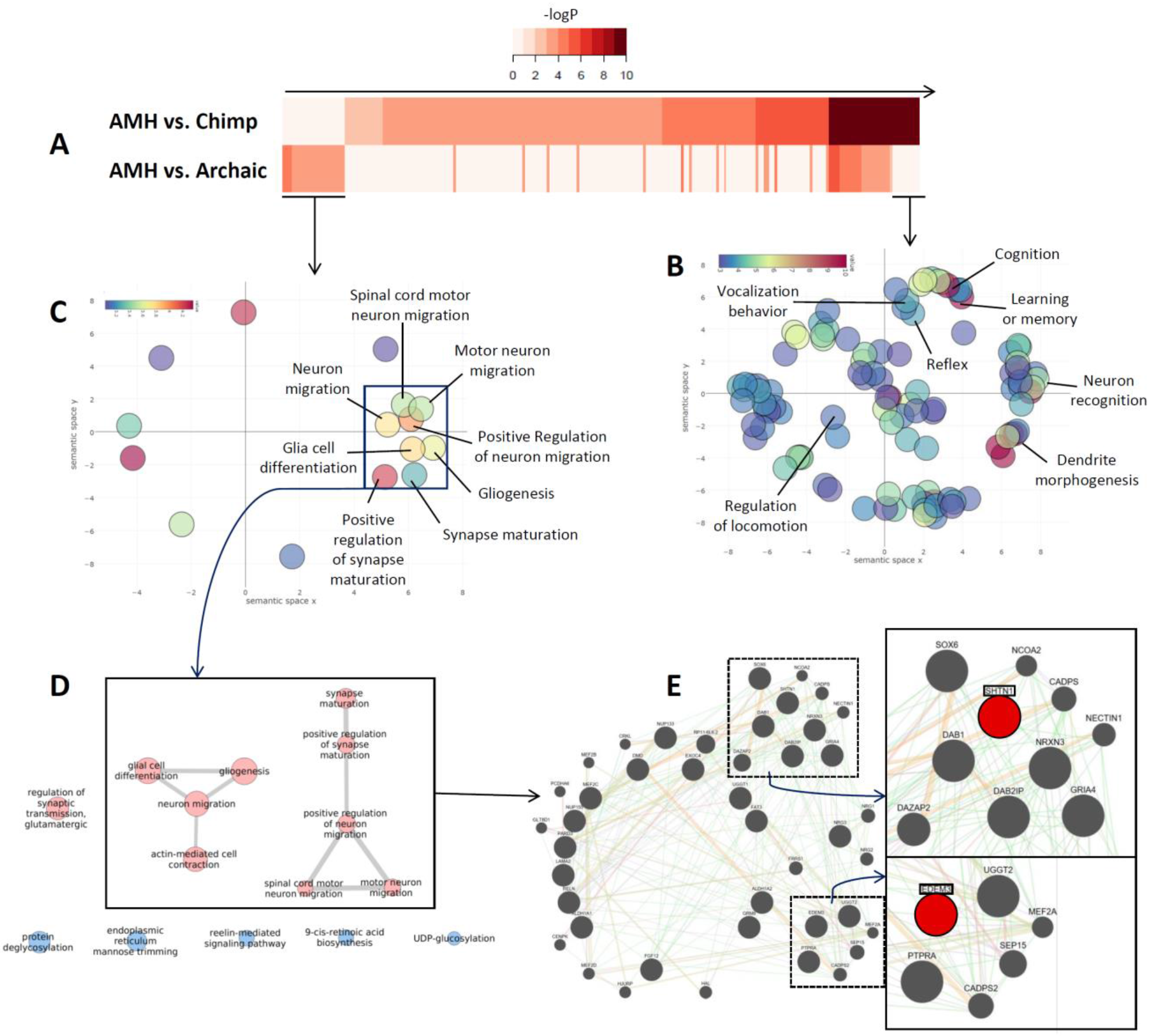
GO analyses of genes targeted by AMH-specific RIs. A) Heat maps representing -log(p-values) of GO terms associated with genes targeted by AMH-specific RIs vs chimp (top) and AMH-specific RIs vs both HN/HD (bottom), order for increased significance in the top row.
B) Scatterplot representation of the identified hominid-specific GO terms. The x and y coordinates of the circles were derived from the Revigo analysis, based on multidimensional scaling on the matrix with the GO semantic similarity values. The functional categories associated with genes that form networks are highlighted and labeled. C) Scatterplot representation of the identified sapiens-specific GO terms. The x and y coordinates of the circles were derived from the Revigo analysis, based on multidimensional scaling on the matrix with the GO semantic similarity values. The functional categories associated with genes that form networks are highlighted and labeled. D) Functionalities of sapiens-specific GO terms associated in networks (if applicable). Red terms are neural-related while blue terms are not. E) Gene network of genes containing AMH-specific RIs absent in both HN and HD with neural functionalities. The larger the circle, the more the gene represented by it has interactions with other genes in the network. The sub-network showing strong interactions with the gene SHTN1 is highlighted in the top-right, while the sub-network with more interactions with the gene EDEM3 is highlighted in the bottom-right.

### Evidences of RI contribution in the molecular differentiation of AMH

Since AMH-specific RIs seem to increase the variability of transcripts and tissue-preferential expression of their targeted genes, we next characterized in greater detail the insertional loci of the three “recent” insertions with a peculiar population distribution that were identified above in paragraph “AMH-specific RIs in present-day populations”. The first one, an AluYa5 RI in chr16q22.1 (Fig. 3A-B), is polymorphic in AFR populations (average frequency of 55%), but fixed or almost fixed in all non-African populations (with the highest difference in frequency between AFR and non-African populations). Although no role in functional alteration was detected for this RI in its insertional locus, the insertion is associated with signs of post Out-of-Africa selection, as revealed by the 3P-CLR selection estimate that places its genomic locus in the top 0.1% loci. The second RI analyzed, an AluYg6 insertion in chr1q25.3, inserted in gene EDEM3 (mentioned above) and is estimated to be fixed or almost fixed in all AMH populations, while completely absent in chimps, HN and HD, as well as in other primates (Figure 3C-D). Remarkably, there is a shorter EDEM3 annotated alternative transcript ending precisely in correspondence with the poly-A tail of the AluYg6 insertion, resulting in exonization of this RI. This alternative EDEM3 transcript is not annotated in chimpanzees, and we suggest that is extremely likely that this transcript is a direct consequence of the AluYg6 insertion into gene EDEM3. Finally, the third RI analyzed, an AluYb9 RI in chr10q25.3, inserted in the 15th intron of the gene SHTN1, which has the highest level of interaction in the previously identified network of neural genes, and the Alu inserted antisense in respect to the gene’s transcriptional directionality (figure 3E-F). This specific RI is mostly fixed in all AMH populations and absent in HN, HD and chimp genomes, as well as in other primates.

**Figure 3.**
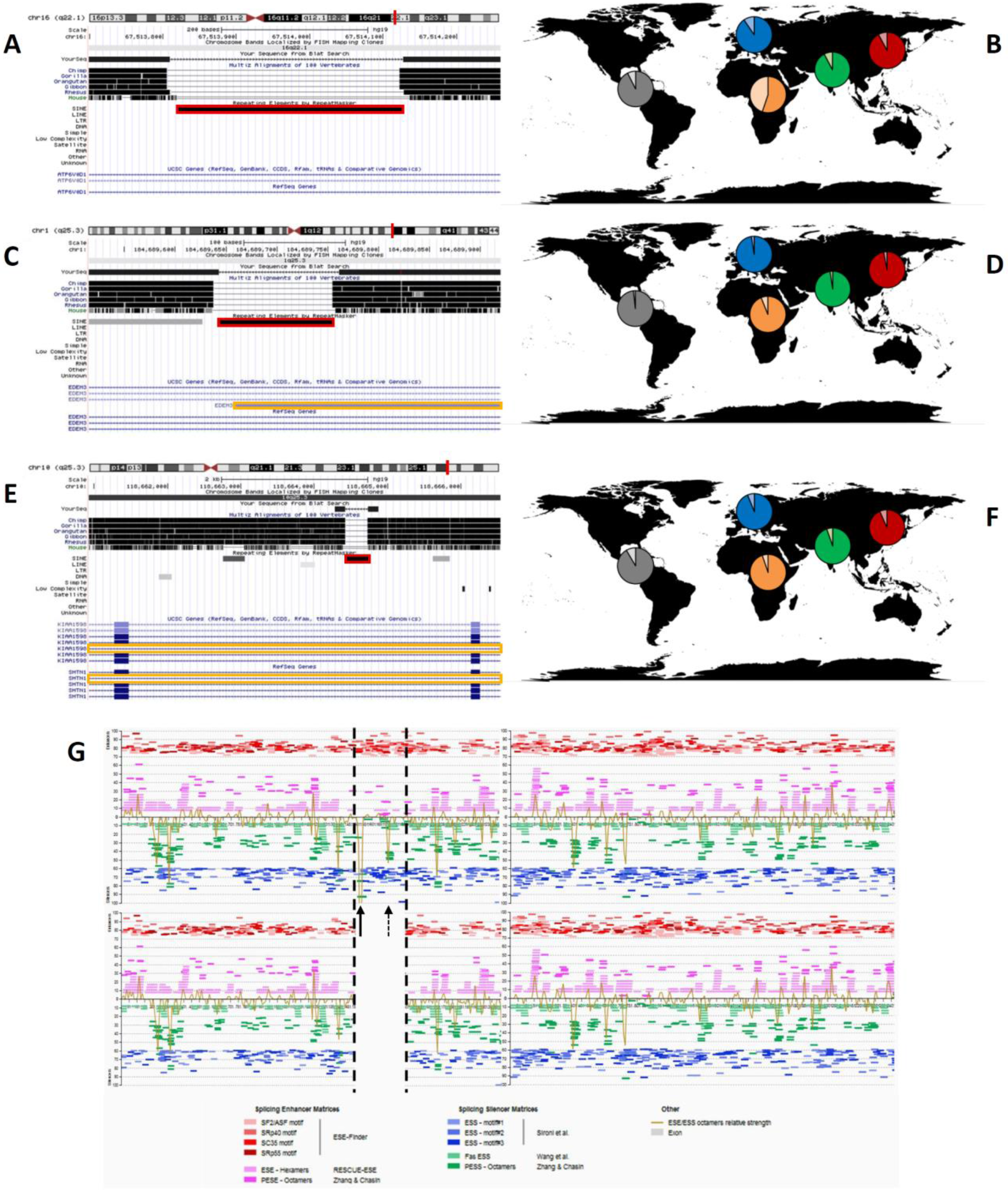
The impact of AMH-specific RIs. A-B) Annotation of the genomic location and distribution in present-day populations of the AluYa5 insertion on chr16q22.1. The insertion is highlighted in red in A; in B, for each diagram, a darker color indicates the presence of the RI and a lighter one its absence. C-D) Annotation of the genomic location and distribution in present-day populations of the AluYg6 insertion on chr1q25.3. In C, the insertion is highlighted in red and a yellow rectangle highlights an alternative transcript that terminates precisely at the poly-A tail of the RI; in D, for each diagram, a darker color indicates the presence of the RI and a lighter one its absence. E-F) Annotation of the genomic location and distribution in present-day populations of the AluYb9 insertion on chr10q25.3. In E, the insertion is highlighted in red and yellow rectangles highlight annotated alternatively-spliced products for the gene in which the insertion occurred; in F, for each diagram, a darker color indicates the presence of the RI and a lighter one its absence. G) Splicing prediction in the sequence corresponding to filed allele (top, containing the intron and the AluYb9 insertion on chr10q25.3) and in the sequence corresponding to the empty allele (bottom, containing just the intron). The sequence is oriented in the same transcriptional sense orientation of the gene, black dotted lines highlight the position of the RI. Pink and red lines represent Splicing Enhancer Matrices, green and blue ones Splicing Silencing Matrices; ochre lines represent the combined strength of Enhancer/Silencing Matrices on the sequence. Arrows highlight silencing signal peaks that occur precisely in the RI sequence.

The gene SHTN1 has various annotated transcriptional/splicing variants, two of which lack the first the two exons that flank the intron where the Alu insertion occurred. Intriguingly, the analyses of this intron with Human Splicing Finder 3.0 (Desmet et al. 2009), both as an empty allele and as a filed allele with the AMH-specific AluYb9 insertion, revealed that differences in the predicted Splicing Enhancing/Silencing Matrices are present between the two sequences. Remarkably, these analyses revealed the presence of putative splicing silencing peaks in the filed allele. The strongest peak is located precisely in the inserted AluYb9 sequence (Figure 3G), suggesting that the Alu insertion induced a splicing-silencing effect on the SHTN1 gene.

## DISCUSSION

In AMH, retrotransposition has been studied mostly for its mutagenic effects and implications in disease insurgence. On the other hand, knowledge about the molecular evolution of our genome relies mostly on simple markers such as SNPs, short InDels and large Copy Number Variants (CNVs), while the role of repetitive/complex regions of the genome is poorly understood. Among others, the complexity of analyses involving repetitive sequences and structural variations in conjunction with the widely used NGS sequencing technology is a challenging task. However, evidences from various Eukaryote organisms suggest that retrotransposition might play an important role in speciation and molecular evolution of genomes (Richardson et al. 2015).

In this study, we evaluated the possible impact of RIs on the differentiation processes that occurred in the human lineage and especially during AMH evolution. In order to do this, we first identified putative species-specific RIs across the genomes of AMH, HN, HD and chimpanzees (Table 1). Notably, the identified RIs reveal a relatively constant retrotransposition/maintenance rate in the human lineage (0.6-0.8 insertions/Ky), which is ^~^2.5x higher than the rate of N-LTRr mobility/maintenance in chimps (0,29 insertions/Ky). In sum, these data suggest that n-LTRr might have, in a constant manner, impacted our genome throughout the evolution of the human lineage more than they have affected the chimp lineage, despite chimpanzees’ shorter generation time. However, the impact of RIs described in this study is just a minor repertoire of the putative effect that RIs can exert on genome structure and regulation, as i) our study is limited to the identification of RIs on very limited sequencing information from Neanderthal, Denisovan and Chimpanzee genomes; and ii) in this study we could only analyze the impact of RIs in *cis*, although RIs are known to impact gene expression and genomic architecture both in *cis* and in *trans* (Garcia-Perez et al. 2016). Thus, we are just starting to uncover the role of n-LTRrs on human evolution, and future genomic studies on Neanderthals and Denisovans will help revealing the full impact of RIs on the evolution of the human lineage.

### AMH-specific RIs and functional variability increase at targeted loci

Focusing on the insertions that are specific to AMH, one of our most prominent results is the strong correlation highlighted between RIs that integrated in genes and the increase in the number of annotated transcript for the genes in which the insertion occurred (Figure 1B-C). While this is true for all RT-DB insertions and the corresponding genes, the effect seems even higher for RIs that are exclusive to AMH. This could be interpreted as a sign of target-preferentiality of retrotransposons in general and particularly in the human lineage. RIs might thus preferentially target genes with a high variety of transcripts. However, the reported random nature of retrotransposition (Richardson et al. 2015), together with the characteristics of N-LTRr sequences and their possible effects upon integration (Goodier and Kazazian, 2008; Cordaux and Batzer, 2009), strongly suggest that the increase in the variety of transcripts is an effect, and not a cause, of the accumulation of new retrotransposition events. It is tempting to speculate that the new transcript/splicing variants of genes targeted by AMH-specific RIs has led to an increase in their functional complexity.

Another important observation from the analyses included in this study is that genes targeted by AMH-specific RIs tend to be preferentially expressed in specific tissues (Figure 1D-E). Notably, in the human lineage these genes are especially likely to be expressed in the brain, with an enrichment of >26% compared to its preferentiality for all human genes. In particular, those genes targeted by AMH-specific insertions vs chimp are more expressed in the amygdala, hippocampus and undifferentiated neurons (up to >52% in respect to each cell-type/tissue expectation), while after the split with HD and HN the enrichment of preferential expression occurs specifically in undifferentiated neurons (>85% in relation to their general baseline). These results show a strong association between RIs and neural genes in the lineage of AMH. Indeed, GO analyses revealed a consistent pattern of neural-related functionalities for genes containing RIs, which is consistent with the aforementioned tissue-specific preferential expression. Interestingly, hominid-specific GO terms of genes targeted by AMH-specific RI vs chimp are highly related to biological and ethological processes that occurred during the differentiation of hominids after the split from chimpanzees, including: neuronal signaling, cognitive capacity, vocalization behavior, reflexes and locomotion regulation (Figure 2B). Strikingly, the most relevant functionalities associated with sapiens-specific GO terms all relate to glia differentiation, synapse maturation and neuron maturation and migration (Figure 2C-D). These functionalities, associated with the preferential expression in undifferentiated neurons of genes that carry AMH-specific RIs absent in both HN and HD, might reflect the importance of these genes in human neural differentiation processes. It is therefore tempting to speculate that the aforementioned increase in transcript variability of specific genes, seemingly induced by RIs, could be tied to the increase in functional complexity of the human brain that occurred all throughout our evolutionary lineage (Hrvoj-Mihic et al. 2013).

### Population distribution of RIs and natural selection

These possible variability-increasing effects of RIs in our lineage and their specific relevance in neural genes should, theoretically, have been subjected to natural selection. Among the identified 321 AMH-specific RIs, the most likely candidates for an adaptive effect on their carriers are those insertions that are fixed across all AMH populations or whose distribution reflects a strong phylogenetic/phylogeographic pattern (e.g. fixation post Out-of-Africa). We therefore compared the identified RIs in this study with the genomic variability of present-day human populations provided by the 1000 Genomes Consortium data. AMH-specific RIs, according to calculated TMRCAs (for AFR populations in which the insertion is fixed or almost fixed), seem to have occurred between >6.5 Mya (i.e. before the split with the chimpanzee lineage) and present day (Supplemental Table S10). These time estimates must, however, only be intended as upper limits for the actual times of the insertions occurrence. Indeed, a large portion of identified putative HN- and HD-specific insertions was shown to be polymorphic to various degrees in present-day populations. Interestingly, the absence of some archaic-specific insertions in all AFR populations and their presence in some (or all) non-African individuals strongly suggest introgression of genetic material in AMH that occurred post Out-of-Africa, possibly via interbreeding with HN and HD. This observation and the populations in which this phenomenon seems to have occurred is consistent with previously reported examples of interbreeding evidences between AMH and HN/HD after *Homo sapiens* migrated into Eurasia (Green et al. 2010; Prüfer et al. 2014; Gardner et al. 2017).

As expected, RIs seem to have been selectively neutral and polymorphic throughout AMH populations although in some instances they show traces of selection only a long time after their putative occurrence (Supplemental Tables S10 and S11). Amongst all RIs that are present at high frequencies in modern populations, a few of them show a peculiar geographic distribution characterized by polymorphism in Africa and fixation (or almost fixation) in non-African individuals. We hypothesize that these RIs could have reached fixation following the Out-of-Africa event as a consequence of genetic drift or selection. In both cases, these insertions predate the spread of AMH out of Africa. In particular, the AluYa5 insertion in chr16q22.1 identified in this study (Figure 3A-B) has a TMRCA of ^~^300 Kya and displays rapid fixation in non-African populations, with its surrounding locus being in the top 0.1% 3P-CLR loci. Therefore, we speculate that this insertion was actually subjected to selection post Out-of-Africa, possibly hundreds of Ky after the insertion itself occurred. While our estimations are only approximations for both the putative age of an insertion and selective pressures acting on an insertional locus, due to the lack of specific methodologies of time estimation and selection for RIs, our results suggest that RIs might occur in a genome and can be maintained randomly within a population under neutral selective pressures. At later times, because of population dynamics or environmental changes, an insertion and the putative novel functional variants it generated might be co-opted and undergo non-neutral selective pressure, in a similar manner as previously reported cases of “soft” selective sweeps detected with SNPs analyses (Pritchard et al. 2010; Hernandez et al. 2011). On an evolutionary timescale, this process seems more likely than insertions having a strong functional-alteration effect immediately upon integration. In fact, most functional regions of a genome are highly conserved and functionality-altering effects would likely be disease-inducing and selected against alleles carrying the RIs. Additionally, genetic drift might also play an important role in the maintenance/diffusion of RIs in human populations, particularly concerning Out-of-Africa bottlenecks.

This interpretation is also consistent with differences in the percentage of RIs targeting genes observed in RT-DB with respect to AMH-specific RIs, both vs chimp and vs HD/HN (Figure 1A). Indeed, these three datasets of retrotransposition events are progressively smaller subsets of the same starting pool of insertions and reflect progressively shorter timescales. Importantly, the effects of a retrotransposon insertion can be co-opted even a long time after the insertion itself occurred, creating new functional variants; thus, a dataset of older insertions (on average) is more likely to show annotated functional variants in a modern genome than a dataset of relatively younger insertions.

### The impact of RIs in modern humans

Previous studies have revealed how retrotransposons can influence the regulation of the loci in which they inserted in a myriad of ways (Richardson et al. 2015). Besides the activity of the sense and antisense LINE-1 promoters contained within full-length LINE-1s (Swergold 1990; Speek 2001; Macia et al. 2011) and the epigenetic silencing of retrotransposon sequences mediated by DNA methylation (Yoder et al. 1997) or histone modifications (Garcia-Perez et al. 2010; Bulut-Karslioglu et al. 2014) that can directly impact gene expression, other common effects of RIs on targeted genes include premature transcript termination (Perepelitsa-Belancio and Deininger 2003; Han et al. 2004) and alternative post-transcriptional processing of genes (Belancio et al. 2006; Heras SR et al. 2014; Goodier and Kazazian, 2008; Cordaux and Batzer, 2009). Some of these functional impacts are generated by RIs inserted in genes because of the A/T richness of the LINE-1 sequence (Han et al. 2004) and due to the presence of a poly-A tail at the 3’ end of the retrotransposon insertion (in both LINEs and SINEs), which can increase the repertoire of transcripts produced from the targeted gene (i.e., generating alternative transcripts). Similarly, Alu elements carry a functional polymarase-III promoter that can directly influence gene expression (Murphy and Baralle 1983); additionally, selected Alu insertions can affect the expression of targeted genes by additional mechanisms (Levanon et al. 2004; Pandey and Mukerji 2011; Elbarbary et al. 2013; Morales-Hernández et al. 2016). Indeed, the AluYg6 insertion on chr1q25.3 identified in this study (Figure 3C-D) seems to directly impact EDEM3 gene expression, as an alternative annotated transcriptional variant in humans terminates precisely in the AluYg6 poly-A tail. Intriguingly, EDEM3 belongs to a group of proteins that accelerate degradation of misfolded or unassembled glycoproteins in the Endoplasmic Reticulum (Hirao et al. 2006). The EDEM3 gene also has a large number of associations in the functional network of neural-related genes containing AMH-specific insertions that are absent in both HN and HD, suggesting a strong relevance for EDEM3 in this network (Figure 2E).

Another important effect that RIs can exert on gene expression, especially antisense insertions in respect to the gene’s transcriptional orientation, is the alteration of the post-transcriptional processing of mRNAs, which can result in alternatively-spliced RNAs (Keren et al. 2010). Indeed, the identified AluYb9 insertion on chr10q25.3 that occurred in the 15^th^ intron of the SHTN1 gene (Figure 3E-F-G), is likely altering the post-transcriptional processing of SHTN1 mRNAs. Although based on computational predictions, the splicing-silencing peaks associated with the allele containing the AluYb9 sequence strongly suggest that the post-transcriptional processing of this gene is affected by the insertion, and we speculate that the AluYb9 sequence is inducing alternative splicing of SHTN1 transcripts. In sum, these data demonstrate how intronic RIs can contribute to the generation of novel functional variants exclusive to AMHs. Additionally, the gene SHTN1 is highly expressed in the human brain and is involved in the generation of internal asymmetric signals required for neuronal polarization and neurite outgrowth (Sapir et al. 2013); it is, as well, the gene with most interactions and relevance in the detected network of neural-related genes targeted by AMH-specific RIs vs both HN and HD (Figure 2E). It is worth mentioning that previous studies demonstrated that the SHTN1 gene has undergone positive selection in AMHs, after the split with HN and HD (Weyer and Paabo, 2015). Accordingly, the AluYb9 insertion’s corresponding genomic locus displays a TMRCA of ^~^560 Kya, which is consistent with possible positive selection and spread after the split between the AMH and HD/HN lineages. Thus, the above-mentioned novel functional variants, likely affected by the RI, might thus have contributed to the establishment of the selective process on this neural gene, which in turn may have affected our species differentiation.

## Conclusions

The results presented in this study suggest that non-LTR retrotransposons mediated processes might have played a more than marginal role in recent human evolution. Their distribution in present-day individuals can be useful as a phylogenetic marker and highlights interactions and population dynamics that occurred after the separation from the chimpanzee lineage. RIs display patterns of maintenance and diffusion in modern populations that reflect slow but constant generation of variability. As the new variants can be co-opted at a later moment, selective pressures could arise resulting in the fixation of those variants. Indeed, non-LTR retrotransposon activity results in an expansion of the genic pool of hominids, and can directly generate new functionalities for human genes and/or their transcripts. RIs are also possibly involved, as well, in the differentiation processes of the human brain and its increase in complexity that took place all throughout the evolution of the human lineage. In some instances, as for the AluYg6 insertion on chr1q25.3 and the AluYb9 insertion on chr10q25.3, the effects of RIs on their target in *cis* might have been key contributors to the molecular differentiation of the AMH genomes. Indeed, the impact of non-LTR retrotransposonsons on human evolution described here likely reflect the tip of a much large iceberg, as our study is limited to a few Neanderthal, Denisovan and Chimpanzee genomes, and because only RI-impacts on *cis* could be analyzed. However, the contribution of non-LTR retrotransposition, whose understanding still needs to be further developed, is starting to shed light on the variety and complexity of RI-driven evolutionary processes that shaped our genome and will continue to influence our evolution in the future.

## METHODS

### RI identification between AMH and HN/HD

The methodology is schematically described in Supplemental Figure S1. Letters in the text correspond to conceptual steps in the scheme. All ancient DNA sequencing reads were considered as single-ended for the purpose of RI/flanking sequences interface identification. The methodology uses well-known bioinformatic tools such as the BLAST+ package (Camacho et al. 2008), ABySS (Simpson et al. 2009), BEDTools (Quinlan and Hall, 2010) and RepeatMasker (Smit et al. 2013-2015), implementing them with custom R or Perl scripts for filtering, conversion and general data management.

Step 1. We retrieved consensus sequences of the most recent non-LTR retrotransposons from RepBase (Bao et al. 2015) (AluYa5, AluYb8, AluYb8a1, AluYb9, AluYb10, AluYb11, AluYk13, LINE-1HS, SVA_A, SVA_B, SVA_C, SVA_D, SVA_E, SVA_F), as well as the genomic material of the species to compare (reference sequence GRCh37-hg19 for AMH, the raw reads of the DNA sequencing for both HN and HD). Specifically, the genomes analyzed in this study are those of two individuals, a Neanderthal and a Denisovan, who lived in the Denisova cave at different times (Meyer et al. 2012; Prüfer et al. 2014). These two genomes were selected for their relatively high coverage and ready availability. The selected retrotransposon sequences were identified in both genomes using blastn (A,B), setting the identity parameter to 95%. This was done in order to allow the identification of retrotransposons diverging as much as 5% from their consensus sequence. Because of the repetitive nature of TEs, each insertion has been associated to its unique genomic target. For AMH, this was done by extending TEs matches by 100 bp in the 3’ direction and in the 5’ direction in the reference sequence (3’ and 5’ flanking sequences). The same could not be done for the archaic DNA, having reads averaging 100 bp as a starting point. Many new retrotransposon insertions are 5’ truncated, thus the length and 5’ end of an insertion is not known beforehand. For this reasons, we implemented a custom R script to select the reads that matched at least 30 bp of the 3’ end of the retrotransposon’s sequence and that had at least 30 bp of flanking sequence in the 3’ direction. In order to take account of differential length of the insertions poly-A tails in respect to the consensus sequences we allowed for 25 bp of margin between insertions and flankings. The sequences of the 3’ ends of insertions with their respective flankings were then compared between the two species using blastn, with identity parameter set to 95% (C). Sequences that were present in one species’ DNA and not in the other were selected as putatively species-specific insertions, thus producing two lists: putative archaic-specific insertions 3’ portions (D) and putative modern-specific insertions 3’ portions (E). Step 2.1. The 3’ flanking portions of the putative-archaic specific insertions were used to identify their respective “empty” (pre-insertional) sites in the AMH genome, aligning them with blastn (identity 95%). The selected 3’ portions of the empty sites were extended in the 5’ direction, thus retrieving the sequence corresponding to the 5’ flankings to the putative archaic-specific insertions. The whole empty sites from the AMH genome (200 bp long) and their 3’ and 5’ portions (both 100 bp) were classified in separate sets, using shared codes for sequences belonging to the same site (F). Then, the 5’ portions of the modern-specific empty sites were identified in the archaic DNA-sequencing reads library, using blastn (identity 95%). We then filtered archaic reads containing a match of at least 30 bp to a modern-specific “empty” site’s 5’ portion and at least 30 bp of non-matching bases 3’ of them. These reads should thus contain both the 5’ flanking site and the 5’ terminal portion of the RI (G). The reads pertaining to the two sets of putative archaic-specific insertions 3’ and 5’ portions were associated to their corresponding modern-specific “empty” sites. This allowed to perform *de novo* assemblies site-by-site using the software ABySS (parameter k set to 40, H,I). Only sequences that were unambiguously assembled for both the 3’ and 5’ portions and that had a clear match for a modern-specific empty site were kept to produce the final sets of confirmed archaic-specific insertions 3’ and 5’ portions, as well as confirmed empty sites from the AMH reference genome (J).

Step 2.2. The putative modern-specific insertions 3’ portions were extended to cover the full insertion as well as 100 bp of flankings in both directions (K). Archaic DNA reads were matched against the 3’ and 5’ flanking sequences using blastn (identity 95%). Reads with at least 30 bp match to both flanking sequences were selected. By doing this, the selected reads spanned the whole empty (pre-insertional) site (L). After associating the archaic reads corresponding to the putative empty sites to their respective modern-specific insertions, *de novo* assembly site-by-site was performed with ABySS (parameter k set to 40, M). Only putative modern-specific insertions whose flankings corresponded to an unambiguously assembled archaic empty site were selected as confirmed AMH-specific insertions (N).

After RI identification, all insertions were validated computationally. Putatively modern-specific RI were selected for having only one matching empty (pre-insertional) site unambiguously assembled from ancient DNA reads. All validated AMH-specific insertions and their absence from the assembled archaic empty sites were verified using RepeatMasker. Putatively archaic-specific insertions were instead selected for having unambiguously assembled both portions of each insertions and matching only one empty (pre-insertional) site in the modern reference genome. All archaic-specific insertions 3’ and 5’ portions were verified with RepeatMasker, as was their absence from the modern-specific empty sites. All archaic-and AMH-specific insertions were also verified for presence of the poly-A tail of the inserted element and TSDs flanking the RI.

### RI identification between AMH and chimpanzee reference sequences and RT-DB

First, we retrieved all RI from element families which are known to have been recently active (all AluJ, AluS and AluY subfamilies, all LINE-1HS and LINE1-PA subfamilies, all SVAs) in the two species reference sequences (GRCh37-hg19 and panTro5) from RepBase (Bao et al. 2015). The 5’ and 3’ flanking regions (100 bp) for all retrieved insertions were aligned using blastn (identity 95%) to the genome of the other species in order to find the respective putative empty (pre-insertional) sites. Two matching sequences (at least 85 bp), in close proximity to each other (less than 50 bp), were selected as a putative “empty” site for each “filled” site. These putative empty sites were then aligned back to the first species DNA using blastn (identity 95%) in order to confirm them as pre-insertional loci. After this procedure, we obtained the insertions specific to the first species (i.e. absent in the second) and vice versa.

RT-DB insertions were retrieved from the human reference sequence GRCh37-hg19 and represent all reference insertions of AluS and AluY subfamilies, LINE-1HS, LINE-1PA2, LINE-1PA3, LINE-1PA4 and all SVAs.

### Archaic-specific insertions in 1KG populations and inter-specific RI estimation

All archaic specific insertions loci were checked in 1000 Genomes Phase3 (The 1000 Genomes Project Consortium 2015) .vcf files for the identification of non-reference variants present in modern day individuals. Archaic RI frequency was then averaged in modern populations according to the 1000 Genomes project annotations.

In order to estimate the amount of RI insertions that are polymorphic between populations we checked for the presence of Archaic RI in one AFR individual, then incrementally added other AFR individuals to the comparison. The rate by which polymorphic insertions were identified produced a curve that reaches a plateau after 20 individual confrontations. Applying this model to AMH insertions results in 554 and 376 non-polymorphic between species AMH insertions (vs HN and HD respectively).

### Assessment of AMH-specific RI in 1KG samples and frequency-based population tree

AMH-specific insertions present in the human reference GRCh37-hg19 may be polymorphic within the broader human population. However, lack of aligned reads spanning the insertion is, in itself, necessary but not sufficient to diagnose the absence of a given insertion within an examined resequenced genome. Even if present, indeed, given the high similarity to other copies of the same transposable element elsewhere in the genome, a given insertion may display no aligned reads due to multiple-mapping filterings. To assess presence/absence of a given insertion we therefore estimated the average coverage of the 1100 bps up and down-stream (“surroundings”) of a putative insertion site and compared it with the coverage of the first and last 10 bps within the RI itself (“interfaces”). We therefore avoided any inference based on the coverage of the “core” inserted sequence, since this may have been affected by the multiple-mapping issues described above. We, instead, reckoned that the first and last 10 bps at the interface between the RI and the surrounding loci could be considered unique enough for the mapping algorithm to see them as a single mapping hit. Based on the reads available from the 1000Genomes Phase3 .bam files (The 1000 Genomes Project Consortium 2015) we then considered as:

- “diploid present” an insertion displaying a coverage >0 at both interface regions and where at least one interface region shows a coverage greater than ½ of the average surrounding coverage;
- “haploid present” an insertion displaying a coverage >0 at both interface regions and where both the interface regions show a coverage smaller than or equal to ½ of the average surrounding coverage;
- “absent” if at least one of the interface regions or the surroundings have zero coverage.

Our assessment approach is conservative with respect to the presence of a given insertion, since it is designed to overestimate absence. We then calculated population frequencies of presence of any given insertion, based on all the individuals available from 1000 Genomes Phase 3 (The 1000 Genomes Project Consortium 2015).

For each RI we calculated the absolute delta frequency per each pair of populations and we averaged it for all the insertions. The obtained matrix of average differences in presence/absence of human specific insertions was used to build a neighbour joining tree using the Ape R package (Paradis et al. 2004).

### TMRCA estimates of genetic regions surrounding AMH-specific RI

The time to the Most Recent Common Ancestor (TMRCA) of each 10kbp regions encompassing a given insertion was estimated as described elsewhere (Inchley et al. 2016) based on 1000 Genomes sequences of AFR samples to avoid potential backwards biases due to the documented Neanderthal introgression in Eurasians (Green et al. 2010). All AFR individuals, and not only carriers of an insertion, were used for this calculations.

### 3P-CLR selection estimates for regions surrounding an insertion

For the sites surrounding AMH-specific insertions we aimed at identifying those that underwent positive selection after the split between Africa and Eurasia but prior to population differentiations within Eurasia. To do so, we used the Three Population Composite Likelihood Ratio (3P-CLR) statistic (Racimo 2016), to look for regions in the EGDP dataset (Pagani et al. 2016) that show evidence of selection that likely occurred shortly after the expansion out of Africa. The 3P-CLR statistic assumes a 3-population tree model with no post-split migration. To ensure that the individuals used in the 3P-CLR analyses represent the most basal split within living Eurasian populations, we used for our EAS population only Chinese and Japanese individuals from the Mainland East and Southeast Asia macro-population. The EUR individuals used were a random subset of the South and West Europe and East and North Europe populations. The AFR outgroup population consisted of the Yoruban individuals from the EGDP dataset (Pagani et al. 2016). Following indications (Racimo 2016), 100 SNPs (with at least 20 SNPs between them) were sampled in each window of length 1cM. Upon completion of the scan, sampled SNPs were grouped into 200kb bins that were assigned the maximum 3P-CLR score of the sampled SNPs in the window. Windows containing an AMH-specific insertion site and falling within the top 99th percentile of scores from this 3P-CLR test were considered to be under selection along the shared Eurasian branch.

### AMH-specific RI, genes and preferential expression

Gene-and transcript-annotation tracks for the human reference genome GRCh37-hg19 were retrieved from ENSEMBL (Aken et al. 2016). RI loci for the different databases were identified in those tracks for information on genes containing RI.

Four gene-tracks (All Genes, genes with RT-DB insertions, genes with AMH-specific vs chimp insertions and genes with AMH-specific vs HN/HD insertions) were thus produced and compared for gene proportions and number of annotated transcripts. Proportion of RI occurred in genes were compared between the tracks and tested with Fisher and binomial tests in R.

The tracks were divided in series containing the number of annotated transcripts for each gene, which were then compared between each other and tested with Wilcoxon and Kolmogorov-Smirnov tests in R. Functional annotation data on genes belonging to the aforementioned four tracks were retrieved from DAVID Bionformatics Resources v6.8 (Huang et al., 2009). For general preferential expression information, nomenclature of tissues belonging to cohesive histological complexes was merged under the categories “Brain”, “Testis”, “Epithelium”, “Placenta”, “Uterus”, “Lung”, “Liver”, “Lymph”, “Kidney”, “Eye”, “Muscle”, “Blood”, “Colon” and “Pancreas”. Only tissues individually called for preferential expression by at least 5% of all human genes were selected for the comparison.

For genes preferentially expressed in the brain, the categories were unpacked into “Brain (general)”, “Undifferentiated Neurons”, “Cerebellum”, “Amygdala”, “Hippocampus”, “Peripheral Nervous System”, “Thalamus”, “Cajal-Retzius Cells”, “Cortex”, “Pituitary”, “Hypothalamus”, “Caudate Nucleus”, “Dendritic Cells”, “Substantia Nigra”, “Subthalamic Nucleus”, “Corpus Callosum”. In this case, only tissues individually called for preferential expression by at least 0.5% of all human genes were selected for the comparison. Tissue-by-tissue comparisons were tested using Fisher and binomial tests in R.

### GO functional analysis

To identify the gene-ontology category of the genes targeted by the AMH-specific RI, both vs chimp and HN/HD, we used ToppCluster (Kaimal et al. 2010), which allows the identification of biological programs using different gene sets to perform contrast and comparative analysis. ToppCluster was set with a false-discovery-rate (FDR) threshold of 0.05 and using “GO: biological process” annotation. The obtained matrix was used to compute the −log10 p-values to obtain significance scores for each functional term. Next, to reduce the redundancy within the GO terms, we used REVIGO (Supek et al. 2011) with parameters set to C=0.7, similarity measure “SimRel” (Schlicker et al. 2006) and using the *Homo sapiens* database. The scatterplots showing the representation of clusters from multidimensional scaling of the semantic similarities of GO terms were obtained with R. We used these plot to identify GO terms related with similar biological functions and the associated genes were used as input for GENEMania (default parameter) (Warde-Farley et al. 2010). The networks obtained from REVIGO were downloaded and visualized with Cytoscape (Christmas et al. 2005).

### Case studies insertional loci annotation and functional inference

The three AMH-specific RI absent in both HN and HD that were identified as recent and displaying peculiar population distribution were manually characterized using the UCSC Genome Browser (Speir et al. 2016), including genomic insertional locus, conservation of the sequence among primates, RepeatMasker presence/absence of repetitive elements, gene-and transcript-annotation.

To identify splicing motifs at the level of the insertion in the gene SHTN-1 we used Human Splicing Finder (HSF 3.0) (Desmet et al. 2009). HSF 3.0 was interrogated with the sequence of the RI + 100bp of flanking regions and with the reconstructed flanking without the RI itself. This was repeated for the whole intron where the insertion occurred and for the reconstructed intron lacking the specific RI, in order to assess its possible effect in splicing-alteration.

## AKNOWLEDGEMENTS

SS is supported by the European Research Council ERC-2011-AdG 295733 grant (Langelin). CT is supported by the University of Padova BIRD171214 fund. LP is supported by the European Union through the European Regional Development Fund (Project No. 2014-2020.4.01.16-0024, MOBTT53 grant). JLGP’s lab is supported by CICE-FEDER-P12-CTS-2256, Plan Nacional de I+D+I 2008-2011 and 2013-2016 (FIS-FEDER-PI14/02152), PCIN-2014-115-ERA-NET NEURON II, the European Research Council (ERC-Consolidator ERC-STG-2012-233764), by an International Early Career Scientist grant from the Howard Hughes Medical Institute (IECS-55007420), by The Wellcome Trust-University of Edinburgh Institutional Strategic Support Fund (ISFF2), and by a private donation by Ms Francisca Serrano (*Trading y Bolsa para Torpes*, Granada, Spain).

## REFERENCES

Aken BL, Achuthan P, Akanni W, Amode MR, Bernsdorff F, Bhai J, Billis K, Carvalho-Silva D, Cummins C, Clapham P et al. 2016. Ensembl 2017. Nucleic Acids Res 45: D635–D642.

Baillie JK, Barnett MW, Upton KR, Gerhardt DJ, Richmond TA, De Sapio F, Brennan P, Rizzu P, Smith S, Fell M et al. 2011. Somatic retrotransposition alters the genetic landscape of the human brain. Nature 479: 534–537.

Bao W, Kojima KK, Kohany O. 2015. Repbase Update, a database of repetitive elements in eukaryotic genomes. Mobile DNA 6: 11.

Belancio VP, Hedges DJ, Deininger P. 2006. LINE-1 RNA splicing and influences on mammalian gene expression. Nucleic Acids Res 34: 1512–21.

Bulut-Karslioglu A, De La Rosa-Velázquez IA, Ramirez F, Barenboim M, Onishi-Seebacher M, Arand J, Galán C, Winter GE, Engist B, Gerle B et al. 2014. Suv39h-dependent H3K9me3 marks intact retrotransposons and silences LINE elements in mouse embryonic stem cells. Mol Cell 55: 277–290.

Camacho C, Coulouris G, Avagyan V, Ma N, Papadopoulos J, Bealer K, Madden TL. 2008. BLAST+: architecture and applications. BMC Bioinformatics 10: 421.

Christmas R, Avila-Campillo I, Bolouri H, Schwikowski B, Anderson M, Kelley R, Landys N, Workman C, Ideker T, Cerami E, Sheridan R et al. 2005. Cytoscape: A Software Environment for Integrated Models of Biomolecular Interaction Networks. Am Assoc Cancer Res Educ Book 2005: 12–16.

Chuong EB, Elde NC, Feschotte C. 2017. Regulatory activities of transposable elements: from conflicts to benefits. Nat Rev Genet 18: 71–86.

Cordaux R, Batzer MA. 2009. The impact of retrotransposons on human genome evolution. Nat Rev Genet 10:691–703.

Desmet FO, Hamroun D, Lalande M, Collod-Béroud G, Claustres M, Béroud C. 2009. Human Splicing Finder: an online bioinformatics tool to predict splicing signals. Nucleic Acids Res 37: 9:67.

Doolittle WF, Sapienza C. 1980. Selfish genes, the phenotype paradigm and genome evolution. Nature 284: 601–603.

Elbarbary RA, Li W, Tian B, Maquat LE. 2013. STAU1 binding 3’ UTR IRAlus complements nuclear retention to protect cells from PKR-mediated translational shutdown. Genes Dev 27: 1495–510.

Evrony GD, Cai X, Lee E, Hills LB, Elhosary PC, Lehmann HS, Parker JJ, Atabay KD, Gilmore EC, Poduri A et al. 2012. Single-neuron sequencing analysis of L1 retrotransposition and somatic mutation in the human brain. Cell 151: 483–496.

Fedoroff NV. 2012. Transposable Elements, Epigenetics, and Genome Evolution. Science 338: 758–767.

Feschotte C. 2008. The contribution of transposable elements to the evolution of regulatory networks. Nat Rev Genet 9: 397–405.

Friedli M, Trono D. 2015. The Developmental Control of Transposable Elements and the Evolution of Higher Species. Ann Rev of Cell and Dev Bio 31: 429–451.

Garcia-Perez JL, Widmann TJ, Adams IR. 2016. The impact of transposable elements on mammalian development. Development 143: 4101–4114.

Garcia-Perez JL, Morell M, Scheys JO, Kulpa DA, Morell S, Carter CC, Hammer GD, Collins KL, O’Shea KS, Menendez P et al. 2010. Epigenetic silencing of engineered L1 retrotransposition events in human embryonic carcinoma cells. Nature 466: 769–773.

Gardner EJ, Lam VK, Harris DN, Chuang NT, Scott EC, Pittard WS, Mills RE, 1000 Genomes Project Consortium, Devine SE. 2017. The Mobile Element Locator Tool (MELT): Population-scale mobile element discovery and biology. Genome Res doi: 10.1101/gr.218032.116.

Gerdes P, Richardson S, Mager D, Faulkner G. 2016. Transposable elements in the mammalian embryo: pioneers surviving through stealth and service. Genome Biol 17: 100.

Goodier JL, Kazazian HH Jr. 2008. Retrotransposons Revisited: The Restraint and Rehabilitation of Parasites. Cell 135(1): 23–35.

Green RE, Krause J, Briggs AW, Maricic T, Stenzel U, Kircher M, Patterson N, Li H, Zhai W, Hsi-Yang Fritz M et al. 2010. A draft sequence of the Neanderthal genome. Science 328: 710–722.

Han JS, Szak ST, Boeke JD. 2004. Transcriptional disruption by the L1 retrotransposon and implications for mammalian transcriptomes. Nature 429: 268–274.

Heras SR, Macias S, Cáceres JF, Garcia-Perez JL. 2014. Control of mammalian retrotransposons by cellular RNA processing activities. Mob Genet Elements 4: e28439.

Hernandez RD, Kelley JL, Elyashiv E, Melton SC, Auton A, McVean G, Sella G, Przeworski M, 1000 Genomes Project. 2011. Classic selective sweeps were rare in recent human evolution. Science 331: 920–924.

Hirao K, Natsuka Y, Tamura T, Wada I, Morito D, Natsuka S, Romero P, Sleno B, Tremblay LO, Herscovics A, et al. 2006. EDEM3, a soluble EDEM homolog, enhances glycoprotein endoplasmic reticulum-associated degradation and mannose trimming. J Biol Chem 281: 9650–9658.

Hrvoj-Mihic B, Bienvenu T, Stefanacci L, Muotri AR, Semendeferi K. 2013. Evolution, development, and plasticity of the human brain: from molecules to bones. Front Hum Neurosci 7: 707.

Huang DW, Sherman BT, Lempicki RA. 2009. Systematic and integrative analysis of large gene lists using DAVID Bioinformatics Resources. Nature Protocols 4: 44–57.

Huang S, Tao X, Yuan S, Zhang Y, Li P, Beilinson HA, Zhang Y, Yu W, Pontarotti P, Escriva H et al. 2016. Discovery of an Active RAG Transposon Illuminates the Origins of V(D)J Recombination. Cell 166: 102–114.

Inchley CE, Larbey CD, Shwan NA, Pagani L, Saag L, Antão T, Jacobs G, Hudjashov G, Metspalu E, Mitt M et al. 2016. Selective sweep on human amylase genes postdates the split with Neanderthals. Sci Rep 6: 37198.

Kaimal V, Bardes EE, Tabar SC, Jegga AG, Aronow BJ. 2010. ToppCluster: a multiple gene list feature analyzer for comparative enrichment clustering and network-based dissection of biological systems. Nucleic Acids Res 38: W96–W102.

Kapitonov V, Jurka J. 2005. RAG1 Core and V(D)J Recombination Signal Sequences Were Derived from Transib Transposons. PLoS Biol 3: e181.

Kapusta A, Kronenberg Z, Lynch VJ, Zhuo X, Ramsay L, Bourque G, Yandell M, Feschotte C. 2013. Transposable elements are major contributors to the origin, diversification and regulation of vertebrate long noncoding RNAs. PLoS Genet 9 doi: 10.1371/journal.pgen.1003470.

Kazazian HH Jr, Wong C, Youssoufian H, Scott AF, Phillips DG, Antonarakis SE. 1988. Haemophilia A resulting from de novo insertion of L1 sequences represents a novel mechanism for mutation in man. Nature 332: 164–166.

Keane TM, Wong K, Adams DJ. 2012. RetroSeq: transposable element discovery from next-generation sequencing data. Bioinformatics 29: 389–390.

Keren H, Lev-Maor G, Ast G. 2010. Alternative splicing and evolution: diversification, exon definition and function. Nat Rev Genet 11: 345–355.

Koonin E, Krupovic M. 2014. Evolution of adaptive immunity from transposable elements combined with innate immune systems. Nat Rev Genet 16: 184–192.

Lander ES, Linton LM, Birren B, Nusbaum C, Zody MC, Baldwin J, Devon K, Dewar K, Doyle M, FitzHugh W et al. 2001. Initial sequencing and analysis of the human genome. Nature 409: 860–921.

Lavialle C, Cornelis G, Dupressoir A, Esnault C, Heidmann O, Vernochet C, Heidmann T. 2013. Paleovirology of ‘syncityns’, retroviral env genes exapted for a role in placentation. Philos Trans R Soc B Biol Sci 368: 20120507 doi: 10.1098/rstb.2012.0507.

Lee E, Iskow R, Yang L, Gokcumen O, Haseley P, Luquette LJ, Lohr JG, Harris CC, Ding L, Wilson RK et al. 2012. Landscape of somatic retrotransposition in human cancers. Science 337: 967–971.

Levanon EY, Eisenberg E, Yelin R, Nemzer S, Hallegger M, Shemesh R, Fligelman ZY, Shoshan A, Pollock SR, Sztybel D et al. 2004. Systematic identification of abundant A-to-I editing sites in the human transcriptome. Nat Biotechnol 22: 1001–5.

Lynch V, Leclerc R, May G, Wagner G. 2011. Transposon-mediated rewiring of gene regulatory networks contributed to the evolution of pregnancy in mammals. Nat Genet 43: 1154–1159.

Macia A, Muñoz-Lopez M, Cortes JL, Hastings RK, Morell S, Lucena-Aguilar G, Marchal JA, Badge RM, Garcia-Perez JL. 2011. Epigenetic control of retrotransposon expression in human embryonic stem cells. Mol Cell Biol 31: 300–316.

Meyer M, Kircher M, Gansauge M, Li H, Racimo F, Mallick S, Schraiber JG, Jay F, Prüfer K, de Filippo C et al. 2012. A high-coverage genome sequence from an archaic Denisovan individual. Science 338: 222–226.

Montinaro F, Busby GB, Pascali VL, Myers S, Hellenthal G, Capelli C. 2015. Unravelling the hidden ancestry of American admixed populations. Nat Commun 6:6596 doi: 10.1038/ncomms7596.

Morales-Hernández A, González-Rico FJ, Román AC, Rico-Leo E, Alvarez-Barrientos A, Sánchez L, Macia Á, Heras SR, García-Pérez JL, Merino JM et al. 2016. Alu retrotransposons promote differentiation of human carcinoma cells through the aryl hydrocarbon receptor. Nucleic Acids Res 44: 4665–83.

Moran JV, DeBernardinis RJ, Kazazian HH Jr. 1999. Exon shuffling by L1 retrotransposition. Science 283: 1530–1534.

Muotri AR, Chu VT, Marchetto MC, Deng V, Moran JV, Gage FH. 2005. Somatic mosaicism in neuronal precursor cells mediated by L1 retrotransposition. Nature 435: 903–910.

Murphy MH, Baralle FE. 1983. Directed semisynthetic point mutational analysis of an RNA polymerase III promoter. Nucleic Acids Res 11: 7695–700.

Notwell J, Chung T, Heavner W, Bejerano G. 2015. A family of transposable elements co-opted into developmental enhancers in the mouse neocortex. Nat Commun 6: 6644.

Ohshima K, Hattori M, Yada T, Gojobori T, Sakaki Y, Okada N. 2003. Whole-genome screening indicates a possible burst of formation of processed pseudogenes and Alu repeats by particular L1 subfamilies in ancestral primates. Genome Biol 4: R74.

Orgel LE, Crick FH. 1980. Selfish DNA: the ultimate parasite. Nature 284: 604–607.

Pagani L, Lawson DJ, Jagoda E, Mörseburg A, Eriksson A, Mitt M, Clemente F, Hudjashov G, DeGiorgio M, Saag L, et al. 2016. Genomic analyses inform on migration events during the peopling of Eurasia. Nature 538: 238–242.

Pandey R, Mukerji M. 2011. From ‘JUNK’ to just unexplored noncoding knowledge: the case of transcribed Alus. Brief Funct Genomics 10: 294–311.

Paradis E, Claude J, Strimmer K. 2004. APE: analyses of phylogenetics and evolution in R language. Bioinformatics 20: 289–290.

Perepelitsa-Belancio V, Deininger P. 2003. RNA truncation by premature polyadenylation attenuates human mobile element activity. Nat Genet 35: 363–366.

Pritchard JK, Pickrell JK, Coop G. 2010. The Genetics of Human Adaptation: Hard Sweeps, Soft Sweeps, and Polygenic Adaptation. Curr Biol 20: R208–R215.

Prüfer K, Racimo F, Patterson N, Jay F, Sankararaman S, Sawyer S, Heinze A, Renaud G, Sudmant PH, de Filippo C, Li H et al. 2014. The complete genome sequence of a Neanderthal from the Altai Mountains. Nature 505: 43–49.

Quinlan AR, Hall IM. 2010. BEDTools: a flexible suite of utilities for comparing genomic features. Bioinformatics 26: 841–842.

Racimo F. 2016. Testing for Ancient Selection Using Cross-population Allele Frequency Differentiation. Genetics 202: 733–50.

Reich D, Green RE, Kircher M, Krause J, Patterson N, Durand EY, Viola B, Briggs AW, Stenzel U, Johnson PLF et al. 2010. Genetic history of an archaic hominin group from Denisova Cave in Siberia. Nature, 468: 1053–1060.

Richardson SR, Doucet AJ, Kopera HC, Moldovan JB, García-Pérez JL, Moran JV. 2015. The Influence of LINE-1 and SINE Retrotransposons on Mammalian Genomes. Microbiol Spectr 3: MDNA3-0061–2014

Sapir T, Levy T, Sakakibara A, Rabinkov A, Miyata T, Reiner O. 2013. Shootin1 acts in concert with KIF20B to promote polarization of migrating neurons. J Neurosci 33: 11932–48.

Sayah DM, Sokolskaja E, Berthoux L, Luban J. 2004. Cyclophilin A retrotransposition into TRIM5 explains owl monkey resistance to HIV-1. Nature 430: 569–573.

Sawyer S, Renaud G, Viola B, Hublinc J, Gansaugea M, Shunkov MV, Dereviankod AP, Prüfer K, Kelso J, Pääbo S et al. 2015. Nuclear and mitochondrial DNA sequences form two Denisovan individuals. Proc Natl Acad Sci 112: 15696–15700.

Schlicker A, Domingues FS, Rahnenführer J, T Lengauer. 2006. A new measure for functional similarity of gene products based on Gene Ontology. BMC Bioinformatics 7: 302.

Simpson JT, Wong K, Jackman SD, Schein JE, Jones SJ, Birol I. 2009. ABySS: a parallel assembler for short read sequence data. Genome Res 19: 1117–23.

Smit AFA, Hubley R, Green P. RepeatMasker Open-4.0. 2013-2015 <http://www.repeatmasker.org>.

Speek M. 2001. Antisense promoter of human L1 retrotransposon drives transcription of adjacent cellular genes. Molecular Cell Biology 21: 1973–1985.

Speir ML, Zweig AS, Rosenbloom KR, Raney BJ, Paten B, Nejad P, Lee BT, Learned K, Karolchik D, Hinrichs AS et al. 2015. The UCSC genome browser database: 2016 update. Nucleic Acids Res 44: D717–D725.

Supek F, Bošnjak M, Škunca N, Šmuc T. 2011. REVIGO summarizes and visualizes long lists of Gene Ontology terms. PLoS ONE 6: e21800.

Swergold GD. 1990. Identification, characterization, and cell specificity of a human LINE-1 promoter. Mol Cell Biol 10: 6718–29.

The 1000 Genomes Project Consortium. 2015. A global reference for human genetic variation, Nature 526: 68–74.

Warde-Farley D, Donaldson SL, Comes O, Zuberi K, Badrawi R, Chao P, Franz M, Grouios C, Kazi F, Tannus Lopes C, et al. 2010. The GeneMANIA prediction server: biological network integration for gene prioritization and predicting gene function. Nucleic Acids Res 38: W214–W220.

Weyer S, Paabo S. 2015. Functional Analyses of Transcription Factor Binding Sites that Differ between Present-Day and Archaic Humans. Mol Biol Evol 33: 316–322.

Wu J, Lee WP, Ward A, Walker JA, Konkel MK, Batzer MA, Marth GT. 2014. Tangram: a comprehensive toolbox for mobile element insertion detection. BMC genomics 15: 795.

Yoder JA, Walsh CP, Bestor TH. 1997. Cytosine methylation and the ecology of intragenomic parasites. Trends Genet 13: 335–340.

